# Intrinsic Network Activity Reflects the Ongoing Experience of Chronic Pain

**DOI:** 10.1101/2021.06.30.450604

**Authors:** Pauline Jahn, Bettina Deak, Astrid Mayr, Anne Stankewitz, Daniel Keeser, Ludovica Griffanti, Viktor Witkovsky, Stephanie Irving, Enrico Schulz

## Abstract

Analyses of intrinsic network activity have been instrumental in revealing cortical processes that are altered in chronic pain patients. However, such studies have not accounted for variable time courses of network activity and subjective pain experience. In a novel approach, we aimed to elucidate how intrinsic functional networks evolve in regard to the fluctuating intensity of the experience of chronic pain.

In a longitudinal study with 156 fMRI sessions, 20 chronic back pain patients and 20 chronic migraine patients were asked to continuously rate the intensity of their endogenous pain. Using group independent component analysis and dual-regression, we extracted the time courses of 100 independent components separately for chronic back pain and chronic migraine. We investigated the relationship between the fluctuation of intrinsic network activity with the time course of subjective pain ratings.

For chronic back pain, we found increased cortical network activity for the salience network and a local pontine network, as well as decreased network activity in the anterior and posterior default mode network for higher pain intensities. Higher pain intensities in chronic migraine were accompanied with lower activity in a prefrontal cortical network.

By taking the perspective of the individual, we focused on the processes that matter for each patient, which are phases of relatively low pain and more straining phases of relatively high pain. The present design of ongoing assessment of the endogenous pain can be a powerful and promising tool to assess the signature of a patient’s endogenous pain encoding over weeks and months.

## Introduction

In recent years neuroimaging studies on pain have largely shifted their focus to the analysis of intrinsic functional networks. These studies have analysed the spatial characteristics of patients’ cortical maps in comparison to healthy control subjects and detected a number of networks that are altered in chronic pain patients (Androulakis et al.,2017; Baliki et al., 2014).

A key region is the default mode network (DMN), where a number of effects have been reported. For example, when comparing patients with chronic back pain (CBP) and patients with chronic regional pain syndrome (CRPS) to healthy controls, Baliki et al. found significant local connectivity differences in brain regions within the DMN. CBP patients showed decreased connectivity to the medial prefrontal cortex and increased connectivity to the precuneus and the right lateral parietal region, as well as to regions outside the DMN, such as decreased connectivity to the anterior cingulate cortex and the left anterior insula/inferior-frontal gyrus (Baliki et al., 2014). For the whole brain DMN weighted map, Loggia et al. found a stronger effect in patients with CBP compared to healthy subjects in the medial prefrontal cortex as well as in the left inferior parietal lobule. Furthermore, the authors revealed a stronger DMN-insula connectivity for the patient group (Loggia et al.,2013). A study on chronic migraine found the opposite effect(Androulakis et al., 2017).

Similarly, the salience network of the brain has received considerable interest due to its higher activity when attending to pain (Kucyi et al., 2013). Kim et al. found a significant cluster of increased functional connectivity in the salience network of patients with CRPS compared with healthy controls (Kim et al., 2018). Likewise, Van Ettinger-Veenstra showed increased connectivity in the salience network in patients with chronic widespread pain when compared to controls, particularly in the left anterior insula/superior temporal gyrus (van Ettinger-Veenstra et al., 2019). Similar to the DMN, chronic migraineurs also showed a contrary result by exhibiting a decreased effect for the salience network (Androulakis et al., 2017).

A recent study showed that the amplitude of low-frequency fluctuations was increased in the post- and precentral gyrus, the supplementary motor area, and the anterior cingulate cortex (ACC) in patients with CBP compared to healthy controls (Zhang et al., 2019). Alshelh et al, on the other hand, showed that patients with chronic orofacial neuropathic pain have significantly reduced amplitude of low-frequency fluctuations in precuneus, posterior cingulate cortex (PCC), medial prefrontal cortex and inferior parietal cortex compared to healthy subjects (Alshelh et al., 2018). Makary et al analysed the power spectral density of nucleus accumbens resting-state activity in subacute and chronic back pain patients and revealed loss of power in the low-frequency band which developed only in the chronic phase of pain, suggesting that altered oscillatory activity in the nucleus accumbens could be a signature of chronic pain (Makary et al., 2020).

However, analyses on maps of intrinsic networks can not fully take into account their ongoing dynamics. Therefore, we aimed to investigate whether the temporal dynamics of the fluctuating strength of functional networks in the brain is related to the subjective experience of endogenous pain in two frequent groups of patients: chronic back pain and chronic migraine.

## Materials and Methods

### Participants

The study included 20 patients diagnosed with chronic back pain (CBP - 16 female; aged 44±13 years) and 20 patients with chronic migraine without aura (CM - 18 female; aged 34±13 years). All participants gave written informed consent. The study was approved by the Ethics Committee of the Medical Department of the Ludwig-Maximilians-Universität München and conducted in conformity with the Declaration of Helsinki.

CBP patients were diagnosed according to the IASP criteria (The International Association for the Study of Pain) (Merskey and Bogduk, 1994), which includes a disease duration of more than 6 months (mean CBP: 10±7 years). All patients were seen in a specialised pain unit. CM patients were diagnosed according to the ICHD-3 (Headache Classification Committee of the International Headache Society (IHS), 2018), defined as a headache occurring on 15 or more days/month for more than 3 months, which, on at least 8 days/month, has the features of migraine headache (mean CM: 15±12 years). All CM patients were seen in a tertiary headache centre.

All patients were permitted to continue their pharmacological treatment at a stable dose (Supplementary Table 1 and 2). The patients did not report any other neurological or psychiatric disorders, or had contraindications for an MRI examination. Patients who had any additional pain were excluded. CBP patients with no pain on the day of the measurement were asked to return on a different day. CM patients were tested on a day with headache but not on a day with a migraine attack. Patients were characterised using the German Pain Questionnaire (Deutscher Schmerzfragebogen)(Casser et al., 2012) and the German-version of the Pain Catastrophising Scale (PCS; Supplementary Table 1 and 2; Sullivan et al., 1995). The pain intensity describes the average pain in the last four weeks from zero to 10 with zero representing no pain and 10 indicating maximum imaginable pain (please note that this scale differs from the one used in the fMRI experiment). The Depression, Anxiety and Stress Scale (DASS) was used to rate depressive, anxiety, and stress symptoms with a cutoff of 10 for depression and stress and 6 for anxiety (Lovibond and Lovibond, 1995). A total PCS score of 30 represents clinically-relevant level of pain catastrophising (Sullivan et al., 1995).

Patients were compensated with 60€ for each session. A total of nine patients were excluded: two patients developed additional pain during the study, the pain ratings of five patients were constantly increasing or decreasing throughout the pain rating experiment, and two patients were unable to comply with study requests. Thirty-six patients were recorded four times across 6 weeks with a gap of at least 2 days (CBP = 9±12 days, CM = 12±19 days) between sessions. Four patients (2 CBP and 2 CM) were recorded three times.

### Experimental procedure

During four separate fMRI recordings, patients rated the intensity of their ongoing pain for 25 minutes using an MRI-compatible potentiometer slider. Two CBP patients and two CM patients underwent three recordings. The scale ranged from zero to 100 in steps of five with zero representing no pain and 100 representing the highest experienced pain (Schulz et al., 2019). On a screen a moving red cursor on a dark grey bar (visual analogue scale, VAS) and a number above (numeric analogue scale, NAS) were shown during the entire functional MRI session. The screen was visible through a mirror mounted on top of the MRI head coil. Patients were asked to look only at the screen, focus on their pain with an emphasis on rising and falling pain. The intensity and the changes of perceived pain had to be indicated as quickly and accurately as possible. To minimise head movement, foams were placed around the head and patients were told to lie as still as possible.

We did not include a healthy control group. Although this group would likely show some fluctuating network activity there is no endogenous pain to relate the fluctuating network activity to. A “correlation” with pain ratings that consists of zeros is statistically not possible. Therefore, a healthy control group using this design is not applicable.

### Data Acquisition

Data were recorded on a 3 tesla MRI scanner (Siemens Magnetom Skyra, Germany) with a 64-channel head coil. Using a multiband sequence (factor 2, T2*-weighted BOLD (blood oxygenation level dependent), images were acquired with the following parameters: number of slices = 46; repetition time/echo time = 1550/30 ms; flip angle = 71°; slice thickness = 3 mm; voxel size = 3×3×3 mm^3^; field of view = 210 mm. 1000 volumes were recorded in 1550 seconds (TR = 1.55 s). Field maps were acquired in each session to control for B0-effects. For each patient, T1-and T2-weighted anatomical MRI images were acquired using the following parameters for T1: repetition time/echo time = 2060/2.17 ms; flip angle = 12°; number of slices = 256; slice thickness = 0.75 mm; field of view = 240 mm, and for T2: repetition time/echo time = 3200/560 ms; flip angle = 120°; number of slices = 256; slice thickness = 0.75 mm; field of view = 240 mm. Head motion did not exceed ±2mm or ±2°.

### Data processing - behavioural data

The rating data were continuously recorded with a variable sampling rate but down-sampled offline at 10 Hz. To remove the same filtering effects from the behavioural data as from the imaging data, we applied a 400 s high-pass filter (see below). For the statistical analysis, the resulting filtered time course was transferred to Matlab (Mathworks, USA, version R2018a) and down-sampled to the sampling frequency of the imaging data (1/1.55 Hz). We shifted the rating vector between −25 s and 35 s in steps of 0.5 s (121 steps). These systematic shifts would account for (a) the unknown delay of the BOLD response and (b) for the unknown timing of cortical processing in reference to the rating: some ongoing cortical processes may influence later changes in pain ratings, other processes are directly related to the rating behaviour, or are influenced by the rating process and are occurring afterwards. We are aware that the variable timing of the BOLD response and the variable timing of the cortical processes are intermingled and would interpret the timing aspects with utmost caution.

To disentangle the distinct aspects of pain intensity (AMP - amplitude) from cortical processes related to the sensing of rising and falling pain, we generated a further vector by computing the ongoing rate of change (SLP - slope, encoded as 1, −1, and 0) in the pain ratings. The rate of change is represented as the slope of the regression of the least squares line across a 3 s time window of the 10 Hz pain rating data. We applied the same shifting (121 steps) as for the amplitude time course. A vector of the absolute slope of pain ratings (aSLP - absolute slope, encoded as 0 and 1), represents periods of motor activity (slider movement), changes of visual input (each slider movement changes the screen), and decision-making (each slider movement prerequisites a decision to move). AMP, SLP, and aSLP vectors were convolved with a haemodynamic response function (HRF) implemented in SPM12 with the following parameters: HRF = spm_hrf(0.1,[6 16 1 1 100 0 32]). To avoid any effects of order (in case of continuously rising pain) in the data, the patients’ rating time courses were required to fluctuate at a relatively constant level. Furthermore, cortical analogues of continuously rising pain would not survive the required high-pass filtering.

To ensure the behavioural task performance of the patients fulfilled these criteria, the ratings of each patient’s pain was measured with a constructed parameter PR defined as follows, with “Δpain_ud_/Δtime” being the slope of the regression line of the unfiltered data (*ud*) and σ_fd_ being the sample standard deviation of the filtered data (*fd*):

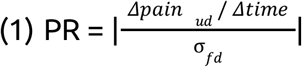

Recordings showing PR values of ≥0.25 were rejected from the analysis or repeated if possible. We excluded five participants and repeated three sessions that exhibited a continuous rise of pain intensity throughout the experiment. The expulsion was caused by premature patient drop-out, as well as by preventing effects of order and the elimination of effects through data filtering.

Taken together, the threshold of the PR value was chosen based on theoretical considerations (effect of order, data filtering), as well as from careful inspection of the data. Subjects with a high PR value are not suitable for a continuous pain rating design in fMRI studies for two reasons. Firstly, steadily changing ratings will cause an effect of order. Secondly, the main changes of brain activity will be removed from the data due to the necessary high-pass filtering.

### Data processing - imaging data

Functional MRI data were preprocessed using FSL (Version 5.0.10), which included removal of non-brain tissue (using brain extraction, BET), slice timing correction, head motion correction, B0 unwarping, spatial smoothing using a Gaussian kernel of FWHM (full width at half maximum) 6 mm, a nonlinear high-pass temporal filtering with a cutoff of 400 s, and linear and non-linear registration to the Montreal Neurological Institute (MNI) standard template. The data were further cleaned of artefacts by performing single-subject ICA with MELODIC. Artefact-related components were evaluated according to their spatial or temporal characteristics and were removed from the data (Griffanti et al., 2014). The average number of artefact components for CM was 40±6 and for CBP 49±8. We deliberately did not include any correction for autocorrelation, neither for the processing of the imaging data nor for the processing of the pain rating time course as this step has the potential to destroy the natural evolution of the processes we aim to investigate (see PALM analysis below).

### Statistical analysis - imaging data

In a next step, we ran a group ICA - separately for CBP and CM with temporally concatenated data of all recordings using MELODIC. The number of components was restricted to 100. Dual regression was then used to derive the corresponding network time series and maps for all components and for each of the sessions. Using Linear Mixed Effects models (LME; MixedModels.jl package in Julia), we aimed to determine the relationship between fluctuating pain intensity and the fluctuating cortical activity separately for each component. The fluctuating network activity of a particular component is modelled through the time course of the three variables (AMP, SLP, aSLP) derived from the pain ratings.

The statistical model is expressed in Wilkinson notation; the included fixed effects (network ~ AMP + SLP + aSLP) describe the magnitudes of the population common intercept and the population common slopes for the relationship between cortical data and pain perception. The added random effects (e.g. AMP - 1 | session) model the specific intercept differences for each recording session (e.g. session specific differences in pain levels or echo-planar image signal intensities):

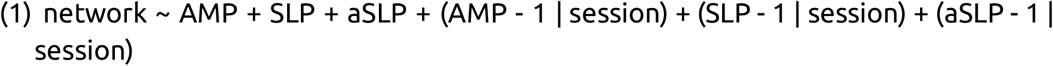

Each model was computed 121 times along the time shift of the rating vector (−25 to 35 s in steps of 0.5 s, see above) and the highest absolute t-values of the fixed effect parameters AMP and SLP were extracted.

### Correcting statistical testing - surrogate data

All statistical tests had to be corrected for multiple testing (components, time shifts) and autocorrelation in the behavioural data: we created 1000 surrogate time courses using the IAAFT algorithm (Iterative Amplitude Adjusted Fourier Transform) from the original rating data, which were uncorrelated to the original rating data but had the same autocorrelation structure as the original data (Schreiber and Schmitz, 1996). Using surrogate data, the entire LME analysis, including the temporal shifts, was repeated 1000 times, resulting in 1000*121*100 statistical tests for AMP and SLP. The highest absolute t-values of each repetition across all components and shifts was extracted. This procedure resulted in a right-skewed distribution of 1000 values for each condition. Based on the distributions of 1000 values (for AMP, SLP), the statistical thresholds were determined using the “palm_datapval.m” function publicly available in PALM (Winkler et al., 2016, 2014).

### Data availability

The data that support the findings of this study are available from the corresponding author upon reasonable request.

## Results

### Questionnaires

The mean pain intensity specified in the questionnaires was 5 ± 2 for CBP and 5 ± 1 for CM (scale ranging from 0 to 10). The mean duration of the chronic pain was 10±7 years for CBP and 15±12 years for CM. The scores for the PCS were 17±10 for CBP and 21±10 for CM. For CBP the depression scale was 4 ± 3, the anxiety scale 3 ± 2 and the stress scale 7 ± 4. For CM the depression scale was 3 ± 3, the anxiety scale 3 ± 4 and the stress scale 6 ± 4 (all results given as mean ± standard deviation). Please see Supplementary Table 1 and 2 for detailed patient characteristics and questionnaire data. A study on healthy subjects found similar results. A large sample of 1794 participants reported scores for depression of 3 ± 4, for anxiety of 2± 3 and for stress of 5 ± 4 (Henry and Crawford, 2005). None of the patients reported any psychiatric comorbidity.

### Behavioural data

The average pain ratings were variable between recording sessions. For CBP and CM we found an average rating of 39 (±14) and 40 (±15), respectively. The pain ratings within each session were substantially fluctuating, as reflected by a high variance over the 25 min of recording: σ^2^=109.3 (±126.6) for CBP and σ^2^=93.3 (±62.8) for CM. In general, we found a minimal positive slope of 0.13 (±0.41; change of rating unit per minute) for CBP and of −0.05 (±0.37) for CM (all mean ± standard deviation). The high similarity of pain ratings is also shown by the largely overlapping within-session distribution of pain ratings for the first and second half of the recording, which amounts to 74±17% for CBP and to 71±13% for CM (area under the curves, AUC). A potential constantly rising pain, which we would have excluded from the present study, would have resulted in 0% overlap.

### Imaging results - Chronic Low Back Pain

For CBP (Figure 1, Table 1) we found two cortical networks whose time course exhibited a significant positive relationship with the time course of pain intensity. We found a positive relation for pain intensity with the salience network, which includes the insular cortex, the frontal operculum, the ACC, the paracingulate cortex (component 12), as well as with a local pontine network (component 60). Further networks, which included the anterior (component 11, aDMN) and the posterior default mode network (component 17, pDMN), exhibited negative relationships. The pDMN includes the angular cortex, the lateral occipital cortex, the precuneus, and the PCC. The pDMN comprises the nucleus accumbens, the nucleus caudatus, the subcallosal cortex, the orbitofrontal cortex, the frontal pole, and the temporal pole.

**Fig. 1.**
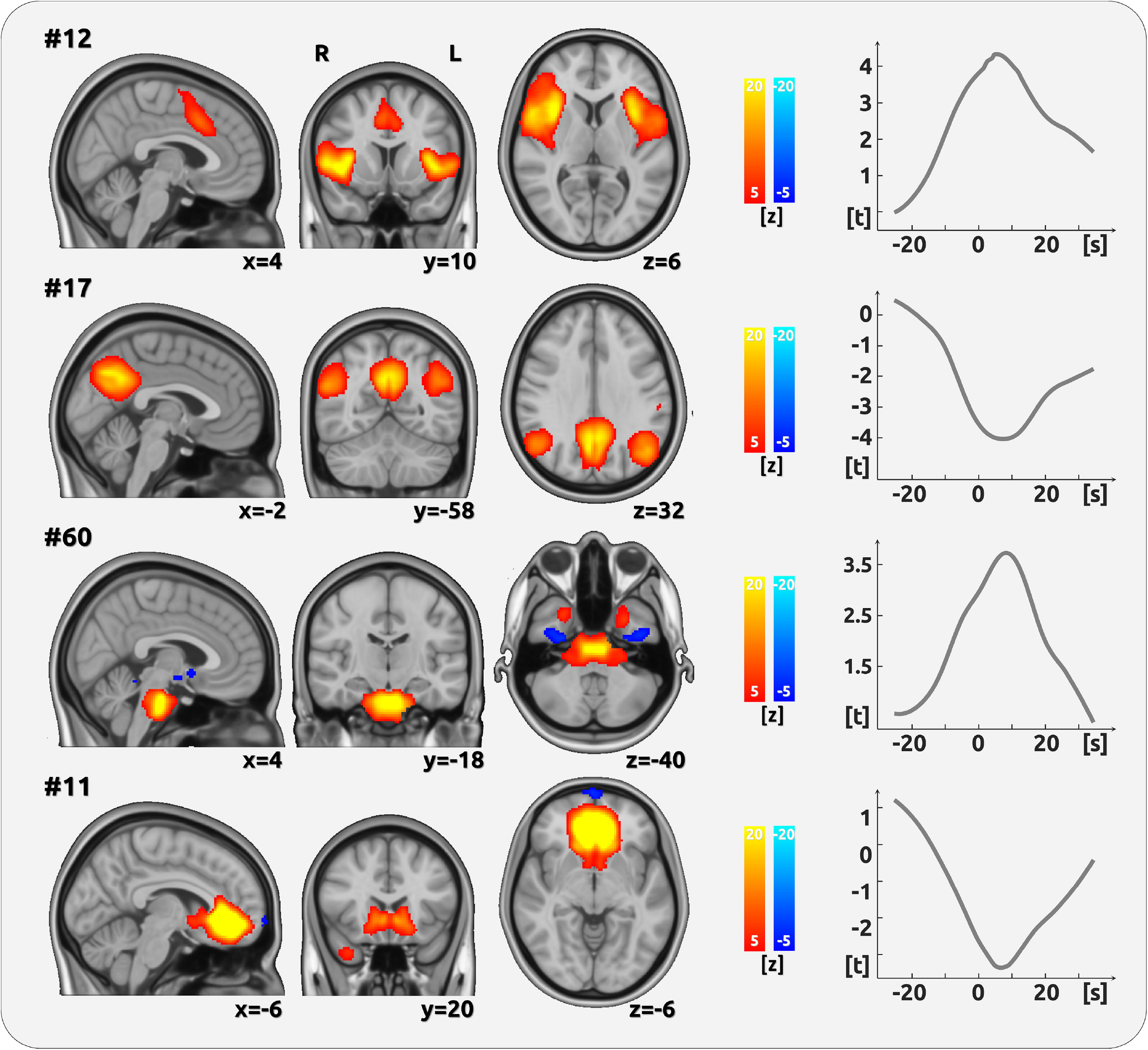
Encoding of pain amplitude for CBP. The figure shows the four networks whose time course exhibited a significant positive relationship with the time course of pain intensity (AMP). The cortical maps on the left side show the spatial characteristics of the networks and are the result of the Melodic ICA. Z-scores were also determined by Melodic and depict the contribution of each voxel to the respective network. The right side exhibits the analysis of the components time courses. The time courses, which are the output from the dual regression, were related to the time course of pain ratings using LMEs. The graph shows the results of the shifts between −25 and 35 s in relation to the current rating (0 s). One significant network component (#12) essentially comprises the insular cortex and the posterior part of the anterior cingulate cortex. A further component with a positive relationship with pain is mainly related to pontine activity (#60). Negative relationships were found for the posterior (#17) and the anterior part of the DMN (#11).

### Imaging results - Chronic Migraine

For CM the time course of activity in a network including the frontal pole, the precuneus and the PCC (prefrontal-precuneus network, component 92 - Figure 2, Table 2) exhibited a significant negative relationship with the time course of pain intensity.

**Fig. 2.**
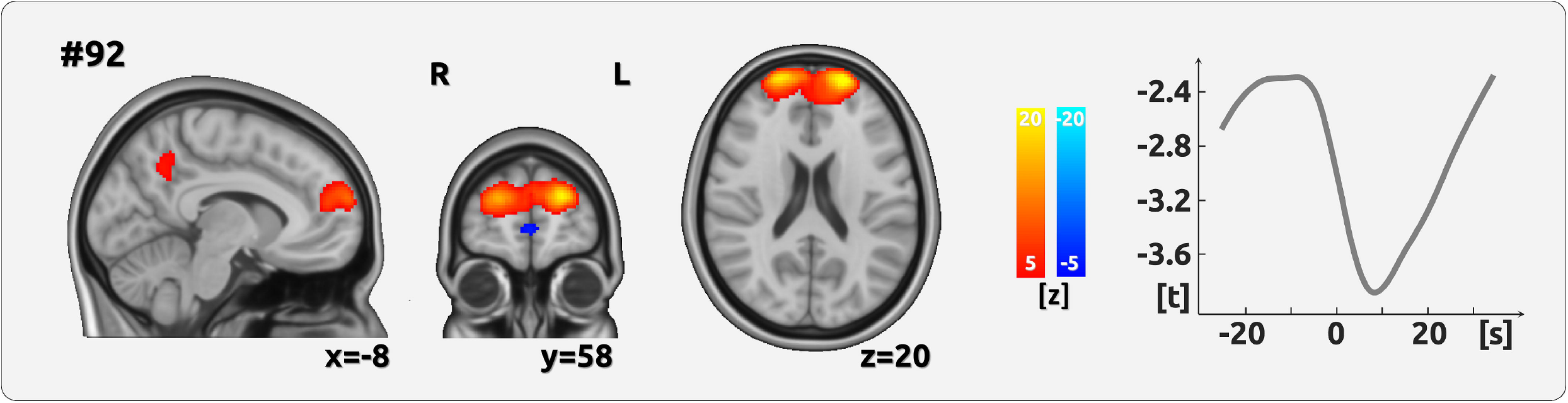
Encoding of pain amplitude for CM. The figure shows the network whose time course exhibited a significant negative relationship with the time course of pain intensity (AMP). The cortical maps on the left side show the spatial characteristics of the networks and are the result of the Melodic ICA. Z-scores were also determined by Melodic and depict the contribution of each voxel to the respective network. The right side exhibits the analysis of the components time courses. The time courses, which are the output from the dual regression, were then related to the time course of pain ratings using LMEs. The graph shows the results of the shifts between −25 and 35 s in relation to the current rating (0 s).The network essentially comprises the prefrontal cortex.

## Discussion

In repeated recordings we investigated in two samples of chronic pain patients how the pain-related intrinsic network activity evolves over time. We did not assume stationary network activity represented by a whole-brain map for each subject that spans the entire testing period, but explored whether the evolving time course of intrinsic networks reflects the fluctuating intensity of the subjective experience of chronic pain.

We revealed variable, non-stationary network activity to represent phases of relatively high and phases of relatively low pain. For higher pain intensities, chronic back pain exhibited an increased network activity for the salience network and a local pontine network, as well as decreased network activity in the anterior and posterior DMN. Higher pain intensities in chronic migraine are accompanied by lower activity in a bilateral prefrontal network. These dynamic aspects of pain can be considered as the core of the patient’s suffering from pain disease. Cortical networks that reflect these dynamic processes may indicate treatment success: patients with higher levels of endogenous pain are relieved by therapies which reduce pain intensity to tolerable levels (Davies et al.,2000; Hirsh et al., 2005).

### Chronic Low Back Pain

#### Default Mode Network

The DMN is thought to represent task-negative experimental periods in the absence of task demands (Li et al., 2020), allowing subjects to fall into a relaxed introspective or self-referencing mode (Schilbach et al., 2012; Sheline et al., 2009). We found a negative relationship between the DMN (anterior and posterior parts) and pain intensity in CBP patients, which may represent the gradual levels a subject can transfer themselves into a certain *relaxed-introspective* mental condition (Kay et al., 2012). Phases of low pain are likely to allow this mode rather than phases of high pain, which may require more attention to the experienced chronic pain input. The DMN has been suggested to be controlled by a network consisting of the anterior insular cortex and the ACC which is largely overlapping with the salience network in the present study (see below) (Li et al., 2018; Sridharan et al., 2008). The DMN has been previously shown to be affected in chronic back pain and other chronic pain states (Alshelh et al., 2018; Baliki et al., 2014). These DMN alterations were interpreted to represent dysfunctions related to a maladaptive physiology (Baliki et al., 2014, 2008). However, the present findings indicate a different interpretation; they are also in line with previous research on the brain-state dependent role of the DMN (Raichle, 2015), reflected in the present study by phases of high (~ high-amplitude DMN) and low pain (~ low-amplitude DMN).

#### Salience network

For chronic low back pain patients, we found a pain-related network that consists of the entire insular cortex and the posterior part of the cingulate cortex. The regions of this networks have - on the one hand - been found in numerous studies on the processing of pain (Davis et al., 2020; Tracey and Mantyh, 2007) and - on the other hand - been interpreted as salience network in resting-state studies (Seeley, 2019; Taylor et al.,2009). In fact, both aspects are not necessarily mutually exclusive (Borsook et al., 2013). The combined activity of both regions could represent the unspecific (salience) aspect that has been suggested to contribute to the processing of several sensory modalities, including pain (Mouraux et al., 2011). Therefore, the present data suggest a similar mechanism for the processing of chronic low back pain intensity, whose primary and common aspect appears to be its saliency.

#### Pontine network

We found a positive relationship between the local pontine network activity and the intensity of CBP. There is less evidence of a pontine contribution to the processing of chronic low back pain so far. This might be caused by the various methodological differences between the present study and previous work and a general neglect of BOLD fluctuations in the white matter (Huang et al., 2020). For the present study, the time course of ventral pontine BOLD activity shows a clear and specific effect at the moment of the pain rating as well as a dropping relationship before and afterwards. The contribution of the (dorsal) pons has been shown in a study that investigated structural changes in chronic back pain patients compared to healthy controls (Schmidt-Wilcke et al., 2006).

### Chronic Migraine

#### Prefrontal-precuneus network

For chronic migraine patients, we found a prefrontal-precuneus network to negatively relate to the ongoing fluctuations of pain intensity. For frontal regions, previous studies found structural changes (DeSouza et al.,2020; Jin et al., 2013; Soheili-Nezhad et al., 2019), as well as decreased functional connectivity (Androulakis et al., 2017) in patients with chronic migraine compared to healthy controls.

The absence of any effect in pain-related areas may reflect the complexity of migraine (Burstein et al., 2015). While the detailed pathophysiological mechanisms for the chronification of migraine are not fully understood (Chong et al., 2019; Filippi and Messina,2020), the current data with multiple recordings can open a window to explore stable and individually specific signatures for each patient.

#### Hypothalamus

We found no significant involvement of the hypothalamus in any of the functional networks. This absence of hypothalamic effect is suggested to be related to two aspects. *First*, we did not find any network that comprises a substantial and distinct portion of the hypothalamus. As the algorithm of the ICA requires large fluctuations of cortical signals, this might not be the case for hypothalamic signals, which exhibit rather small and noise-sensitive amplitude fluctuations. *Second*, the hypothalamus was found to be critically involved in episodic migraineurs during the headache (Denuelle et al., 2007) as well as during the premonitory phase (Maniyar et al., 2014), or in chronic migraine patients in response to experimentally applied pain (Schulte et al., 2017). In contrast, we were exploring the fluctuating endogenous head pain intensity in chronic migraineurs outside an attack. Therefore, due to its involvement in cyclic aspects of migraine, the hypothalamus is believed to serve as an attack trigger rather than to encode the intensity of head pain (May and Burstein, 2019). Therefore, the effects of previous studies are not directly comparable with the present finding due to methodological differences, i.e. subjective experience of endogenous pain vs. experimentally applied pain events, pain encoding vs. pain contrasted to baseline, and intrinsic network activity vs. event-related design.

### Fluctuations of network activity in chronic pain

Previous studies investigating intrinsic networks found chronic pain patients had impairments in the DMN (Baliki et al., 2014; Loggia et al., 2013; Napadow et al., 2010) and the salience network (Kim et al., 2018; Kucyi et al., 2013; van Ettinger-Veenstra et al., 2019) when compared with healthy controls. Specifically for chronic migraine, decreased intrinsic network connectivity has also been found for the frontal executive control network (Androulakis et al., 2017). Common to all of these studies is the voxel-wise analysis on individual maps of intrinsic components. These maps are interpreted as representing the *stationary* strengths of functional connectivity across the entire recording period (Buckner et al., 2013). While voxel-wise analyses allow to evaluate local changes in functional connectivity within a network, significant effects in voxels outside of the main regions of the network are often difficult to interpret. Moreover, a mere focus on cortical maps leaves open the question whether there are different physiological or cognitive states during peaks and troughs of the network time series. Therefore, the stationary network analyses may not represent the core pain experience of the patients, which is characterised by its fluctuating amplitude of pain intensity, unpleasantness, and distress.

The present study is focussed on a rather neglected aspect of network activity, which is the *non-stationary* and fluctuating time course of intrinsic network activity. This time course is dominated by the voxels in the centre of the component maps due to their higher statistical weights. Here, we are able to analyse and interpret the peaks and troughs of the ongoing intrinsic network activity by relating it to the ongoing and variable subjective experience of pain. Although we are analysing intrinsic networks, this study is no resting-state study.

### Summary and outlook

The present investigation has targeted how the ongoing perception of chronic pain is subserved in the human brain. By taking into account the perspective of the individual, we focussed on the processes that matter for each patient, which are phases of relatively low pain and more straining phases of relatively high pain. We pursued a group-wise statistical approach in order to reliably assess the encoding of chronic pain in CM and CBP.

The present investigation also permits a glance into potential future investigations. Although we could not take any individual variations into consideration, there are current advances to examine individual trajectories of chronic pain experiences. These trajectories can be unique to the individual and the present design of ongoing assessment of the endogenous pain is a powerful and promising tool for these analyses. With an accompanying and repeated assessment of a patient’s endogenous pain encoding over weeks or months, future therapies could track a patient’s current cortical pattern of pain encoding and relate changes of these patterns to the success of therapeutic interventions. A tailored medical treatment that is based on the changes of a patient’s endogenous pain signature is a promising outlook into the future of medicine.

