## Supplementary Table 1 and 2 for "Intrinsic Network Activity Reflects the Ongoing Experience of Chronic Pain"

**Supplementary Table 2 Characteristics of CM patients reported in questionnaires**

| patient | m/f | age (years) | pain duration (years) | pain medication | pain intensity | PCS | d/a/s |
|---|---|---|---|---|---|---|---|
| 1 | f | 61 | 50 | Sumatriptan 100mg (20x/month) | 7 | 15 | 0/10/9 |
| 2 | f | 27 | 7 | Metamizole 500mg (2-3x/month), Sumatriptan 50mg (1x/month) | 4 | 5 | 0/0/2 |
| 3 | f | 50 | 35 | Sumatriptan 100mg (5-7x/month) | 4 | 24 | 5/0/6 |
| 4 | f | 27 | 8 | Zolmitriptan 20mg (2x/month), Ibuprofen 600mg (7x/month) | 7 | 37 | 1/1/8 |
| 5 | m | 49 | 30 | Ibuprofen 600mg (7-8x/month), Metamizole 500mg (3-4x/month), Paracetamol 500mg (5-6x/month) | 4 | 3 | 0/6/1 |
| 6 | f | 52 | 30 | Ibuprofen 400mg (6x/month), Paracetamol 1000mg (2x/month) | 5 | 11 | 5/10/12 |
| 7 | f | 32 | 15 | Zolmitriptan 5mg (8x/month), Naproxen 500mg (15x/month), Acetylsalicylic Acid( ASA) 250mg (4x/month), Paracetamol 200mg (4x/month), Caffeine 50mg (4x/month) | 4 | 31 | 5/7/7 |
| 8 | f | 21 | 7 | Sumatriptan 50mg (1x/month) | 4 | 10 | 2/1/0 |
| 9 | f | 19 | 7 | none | 4 | 35 | 10/2/6 |
| 10 | f | 46 | 13 | Ibuprofen 800mg (8-10x/month) | 6 | 31 | 7/10/15 |
| 11 | f | 27 | 13 | Triptan (2-3x/month), ASA 250mg (20-25x/month), Paracetamol 250mg (20-25x/month), Caffeine 50mg (20-25x/month) | 4 | 13 | 6/1/10 |
| 12 | m | 53 | 15 | Ibuprofen 600mg (10x/month) | 6 | 20 | 7/2/4 |
| 13 | f | 30 | 6 | Ibuprofen 400mg (4x/month) | 4 | 24 | 1/1/5 |
| 14 | f | 21 | 7 | Ibuprofen 600mg (4-8x/month), Paracetamol 500mg (4x/month), Zolmitriptan 5mg (1-2x/month) | 3 | 15 | 2/0/5 |
| 15 | f | 23 | 8 | Ibuprofen 600mg (2-3x/month) | 7 | 24 | 2/0/3 |
| 16 | f | 28 | 7 | Ibuprofen 500mg (10x/month) | 7 | 11 | 0/1/1 |
| 17 | f | 25 | 5 | Ibuprofen 600mg (5x/month), Zolmitriptan 5mg (1x/month) | 4 | 21 | 5/5/9 |
| 18 | f | 33 | 20 | Paracetamol 500mg (3x/month) | 5 | 32 | 4/0/4 |
| 19 | f | 21 | 9 | Ibuprofen 400mg (6-10x/month), Rizatriptan 10mg (2x/month) | 5 | 30 | 1/1/2 |

| | | | | | | | |
|---|---|---|---|---|---|---|---|
| 20 | f | 43 | 10 | Ibuprofen 400mg (20x/month), Paracetamol 325 mg (8-10x/month), Naproxen 100 mg (8-10x/month), Caffeine 50 mg (8-10x/month), Drotaverine hydrochloride 40 mg (8-10x/month), Pheniramine 10 mg (8-10x/month) | 4 | 22 | 2/3/9 |

m/f: male/female; PCS: pain catastrophizing scale; d/a/s: depression/anxiety/stress
