## Supplementary Table 1 and 2 for "Intrinsic Network Activity Reflects the Ongoing Experience of Chronic Pain"

**Supplementary Table 1 Characteristics of CBP patients reported in questionnaires**

| patient | m/f | age (years) | pain duration (years) | pain location | pain medication | pain intensity | PCS | d/a/s |
|---|---|---|---|---|---|---|---|---|
| 1 | f | 52 | 32 | thoracic | none | 4 | 14 | 1/4/11 |
| 2 | m | 64 | 2 | lumbar | none | 3 | 22 | 3/5/7 |
| 3 | f | 39 | 8 | lumbar | Ibuprofen 600 mg (10-12x/month)) | 4 | 32 | 7/9/16 |
| 4 | f | 41 | 18 | lumbar | Fluoxetine 20 mg (daily), Paracetamol 500 mg (2-3x/month), Orphenadrine 100mg (3x/month) | 3 | 33 | 6/7/11 |
| 5 | f | 60 | 14 | thoracic | Diclofenac 69,82 mg (1x/month) | 4 | 14 | 2/5/6 |
| 6 | f | 39 | 3 | lumbar | Paracetamol 500mg (2-3x/month), Ibuprofen 400mg (2x/month) | 4 | 13 | 2/1/1 |
| 7 | f | 48 | 5 | cervical | Ibuprofen 400mg (4x/month) | 5 | 12 | 4/0/6 |
| 8 | f | 52 | 11 | thoracic & lumbar | Ibuprofen 400mg (5x/month), Metamizole 500mg (2x/month) | 4 | 7 | 1/2/5 |
| 9 | f | 31 | 6 | lumbar | Ibuprofen 800mg (2-3x/month), Metamizole 1000mg (2-3x/month) | 5 | 1 | 0/0/1 |
| 10 | m | 26 | 4 | lumbar | none | 5 | 9 | 0/3/3 |
| 11 | f | 55 | 16 | thoracic & lumbar | Cannabis drops (15x/month) | 10 | 11 | 5/2/6 |
| 12 | m | 65 | 8 | lumbar | Ibuprofen 600mg (10-15x/month) | 4 | 10 | 2/2/3 |
| 13 | f | 31 | 1 | cervical & thoracic | Ibuprofen 400mg (18x/month) | 3 | 25 | 9/6/14 |
| 14 | m | 32 | 7 | thoracic and lumbar | Ibuprofen 600mg (6x/month), Tramadol 100mg (10x/month) | 5 | 22 | 7/3/5 |
| 15 | f | 26 | 10 | lumbar | Ibuprofen 400mg (1x/month) | 4 | 18 | 6/3/8 |
| 16 | f | 55 | 16 | cervical & thoracic | Metamizole 500mg (2-3x/month) | 5 | 2 | 4/0/4 |
| 17 | f | 56 | 15 | lumbar | none | 7 | 36 | 4/4/13 |
| 18 | f | 42 | 11 | thoracic & lumbar | none | 5 | 10 | 1/2/6 |
| 19 | f | 30 | 3 | thoracic | Ibuprofen 400mg (3x/month) | 4 | 27 | 2/4/8 |
| 20 | f | 43 | 10 | cervical & lumbar | Ibuprofen 400mg (5x/month) | 7 | 22 | 5/4/6 |

m/f: male/female; PCS: pain catastrophizing scale; d/a/s: depression/anxiety/stress. The cutoff for depression and stress is 10 and for anxiety 6.
